# Kinetic measurements of the fluorescent protein synthesis facilitate determination of the miRNA activity

**DOI:** 10.64898/2025.12.17.694323

**Authors:** Philipp Andreevich Malakhov, Yuzhe Wang, Wenyu Xue, Zain Nofal, Nikolay Sergeevich Ilyinsky, Anna Smirnova, Roman Nikolaevich Chuprov-Netochin, Margarita Pustovalova, Sergey Leonov, Julian Markovich Rozenberg

## Abstract

Oncogenesis is inevitably associated with microRNA expression deregulation. Thus, development of both miRNA targeting substances or small RNA for ant-cancer therapy have been reported. Specifically, repression of the miR-16-1-3p and miR-16-2-3p activities play pivotal roles in osteosarcoma and many other cancers. The majority of miRNA sensors use protein degradation to measure miRNA activities. Here we report miRNA sensors that use fluorescent protein synthesis rather than degradation to measure miRNA activity. Specifically, miR-16-1-3p and miR-16-2-3p sensors consist of the bidirectional tet-On system driving the expression of the Katusha2S protein that is regulated by the RNA interference and GFP as a reference. These sensors specifically detect mir16-1-3p and mir16-2-3p small RNA mimics in the osteosarcoma cell line after doxycycline induction. Kinetic measurements of the reporter responses to the miRNA mimics revealed that pre-induced sensors reach significant differences from the control faster, within 2h than the sensors that were induced after mimic transfection. Thus, kinetic measurements of the fluorescent protein synthesis during doxycycline induction of the tet-On system are feasible for determination of the small RNA activity.

## Introduction

Osteosarcoma is a deadly cancer with a 5-year survival rate estimated at 50%-60% (1,2). A rapid progression is observed in the 10-30% of the primary cases with metastasis (3), necessitating novel approaches towards osteosarcoma treatment (4). Multiple studies demonstrated pivotal roles of the miRNAs in the osteosarcoma pathogenesis (5–10). Specifically, mir-16 represses development of osteosarcoma primary tumors and metastasis (11–13). Notably, mir-16 represses the development of breast and colorectal cancers as well (14–16) by forming a feedback loop with p53 and regulating the cell cycle genes (17–19). In our previous investigation, we reported that in addition to the leading strand mir-16-5p of the mir-16, two passenger strands: miR-16-1-3p and miR-16-2-3p originating from two different DNA sites have profound anti-cancerous effects on the growth and metastasis of osteosarcoma (11). While the anti-cancerogenic role of miRNAs including mir-16 in osteosarcoma is well established (6,9–13,20,21), delivery of drugs, including miRNA, to the cancer sites, especially the sites of metastasis is challenging (22). Moreover, the cells of the osteosarcoma microenvironment actively consume, transport and exchange endogenous and likely exogenous miRNA influencing mechanisms of drug action and disease progression (23–25). Therefore, to understand the mechanisms of potential drugs, including miRNA mimics, it is not sufficient to use the delivery sensors such as fluorescent protein co-delivered with miRNA at the tumor sites, we need to be able to measure the activity of miRNAs in the cancer or other specific cell types.

A common strategy to detect miRNA activity is to transduce cancer cells with a fluorescent protein or luciferase followed by several copies of miRNA recognition sequence (26–28). In this scenario, miRNA represses expression of the sensor proteins by the Argonaute-2 mediated mRNA cleavage. It was demonstrated that the degradation kinetics of the fluorescent protein determines the responsiveness of the sensors to the miRNA whose kinetics is relatively rapid (29).

One of the methods for detecting miRNA activity is the use of reporter proteins regulated by a tetracycline inducible bidirectional promoter that is sensitive to a tetracycline activator or repressor. In this method one of the proteins is made sensitive to miRNA by the miRNA complementary sequence inserted in 3’-UTR, while the other is used as a reference (30–32).

Mukherii et all reported that repression of the reporter protein depends on the number of repeats of the miRNA sensor element and on the degree of miRNA complementarity to the sensor (30). However, they did not measure the doxycycline activation kinetics.

Haefilger et. all compared specific and non-specific effects of miRNA but the kinetics of activation was not reported (31). The Tet-On reporter system with a reference protein that is not sensitive to the doxycycline was sensitive to mir-20 sponging but the kinetics of activation in response to doxycycline was not reported (32).

However, it is possible to measure miRNA activity based not on the kinetics of protein degradation, but based on the protein synthesis. If the reporter is induced and then the exogenous miRNA is introduced to the system, then the level of transcript we seek to inhibit is high. In addition, the rate of protein degradation, which is 1.5h at best, determines the rate at which the reporter responds to the changes of miRNA activity (29). In contrast, if we first introduce miRNA into cells and then turn the reporter on, then the transcript is increasing from the zero level and the rate of protein maturation for the fluorescent proteins is about 15 minutes, determining fast response to the changes of the reporter transcript. Therefore, it is possible that this method is more sensitive, however, it was used only for the luciferase reporter (33) and most of the studies are performed on the constitutively induced systems (30–32).

Therefore, we made a bidirectional tet-On system based on Katushka 2S, enabling in vivo detection (34), followed by two copies of the mir16-1-3p or mir16-2-3p complementary sequence (29) and GFP protein transcribed in the opposite direction as a reference.

Moreover, while sensors of the mir-16-5p leading strand are described in the literature (27), we are interested in measuring specific activities of the mir-16 passenger strands: mir-16-1-3p and mir-16-2-3p, that have a profound effects against osteosarcoma (11,12).

We propose that the kinetic measurements of miRNA reporter activity during doxycycline induction that are based on the protein synthesis kinetics will allow fast measurements of the miRNA activity.

To check this hypothesis, we compared the kinetics of the reporter that was pre-activated by doxycycline for 20 h with the reporter that was activated 4h after the exogenous miRNA mimics delivery.

## Methods

### Reporter Cloning

The reporter (Figure 1) was constructed based on a lentiviral plasmid encoding GFP under the control of a tetracycline promoter, as well as expressing a tetracycline activator and containing a puromycin-selective antibiotic (addgene #162823, the gift of Victor Tatarsky) (35). Katushka2S was amplified from the FP761 plasmid from Evrogen. The final construct was assembled using PCR with Q5 polymerase (New England Biolabs) and cloned using XL10-Gold, which ligates DNA at the homologous ends (35). Recognition sites for restriction enzymes were also created around each element in the final construct to facilitate further modifications. In the control plasmid there is a spacer between the MluI and EcoRI restriction sites downstreat Katushka2S that was replaced by two miRNA complementary sequences for mir16-1-3p or mir16-2-3p (Table 1) (29).

**Table 1.**
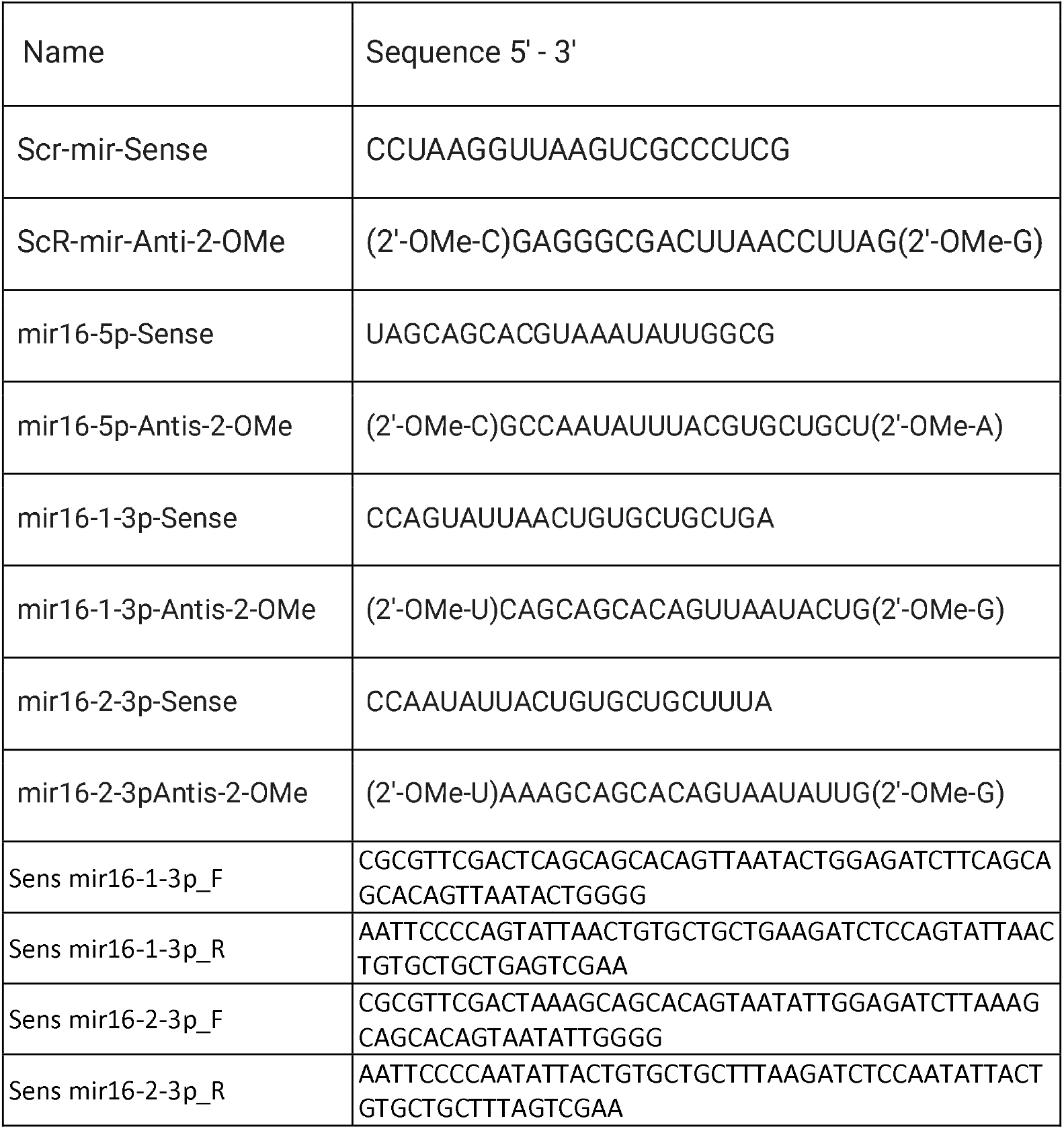
Sequences of miRNA mimics and miRNA sensors.

**Figure 1.**
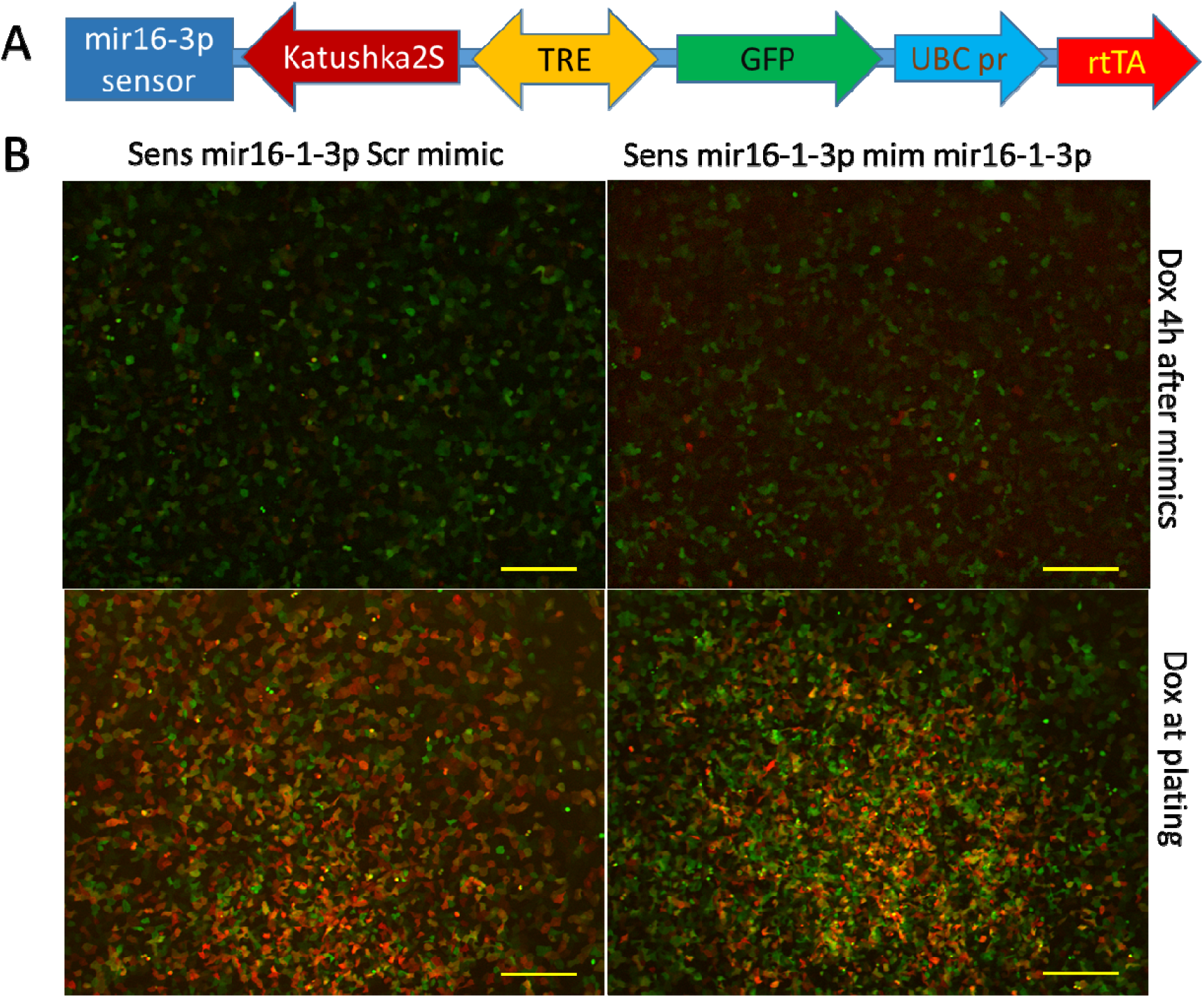
Scheme of the miRNA sensors. Cellular fluorescence was measured 14 and 38 hours after doxycycline induction with the mimics mim Scr and mir16-1-3p over an 18-hour period.

Lentiviruses were produced using standard methods (37). Sequencing and primer synthesis were carried out by Syntol.

### Cell Lines

The U2OS osteosarcoma cell line was transduced by the corresponding lentiviruses, and three days later was followed by the puromycin selection at a concentration of 10 µg/ml for one week. The obtained constructs had up to 85% GFP-positive cells at a high concentration of doxycycline.

We compared two experimental conditions: first - cells that were induced by doxycycline after seeding and 20h later transfected with the mir16-1-3p, mir16-2-3p and scrambled mimics, and second - cells that were transfected with the mimics followed by the doxycycline induction 4h later. The mir16-1-3p and mir16-2-3p mimics consisted of RNA strands paired with corresponding complementary RNA sequences that were 2-O-methylated at the ends (Table 1).

Twenty hours prior to transfection, 40,000 U2OS cells expressing sensors were seeded in a 96-well plate, and 0.005 g/L doxycycline was added to half of the wells. Transfection was performed using 0.3-0.4 µL of GenJet-40 (a gift from Molekta) and 1 µg of mimetics in 100 µL of medium per well of the 96-well plate. Four hours after transfection, the medium was replaced with fresh medium containing doxycycline, and the doxycycline was also added to the wells that did not have doxycycline at seeding.

### Microscopy and Flow Cytometry

Live microscopy was started four hours after transfection at 4x magnification using a JuliaStage microscope, with images taken every two hours or, in some experiments, every hour according to the manufacturer’s recommendations. After data collection from the microscopy, the cells were trypsinized for flow cytometry that was completed within 3 hours.

## Results

### The mir16-1-3p and mir16-2-3p miRNA sensors are specifically inhibited by the corresponding miRNA mimics

The microscopic images revealed that 14 and 38 hours after doxycycline induction, fluorescence was visible (Figure 1). Apparently, the fluorescence was brighter after 38 hours and was somewhat lower in the mir16-1-3p cells in the presence of the corresponding mir16-1-3p mimic (Figure 1).

Approximately 70% of cells were GFP positive by flow cytometry 48 hours after doxycycline induction (Figure 2). Among these, about 25% to 30% of the cells were positive for GFP but did not express Katushka 2S (Figure 2). Conversely, there were relatively few cells that were positive only for Katushka 2S (Figure 2 A, B).

**Figure 2.**
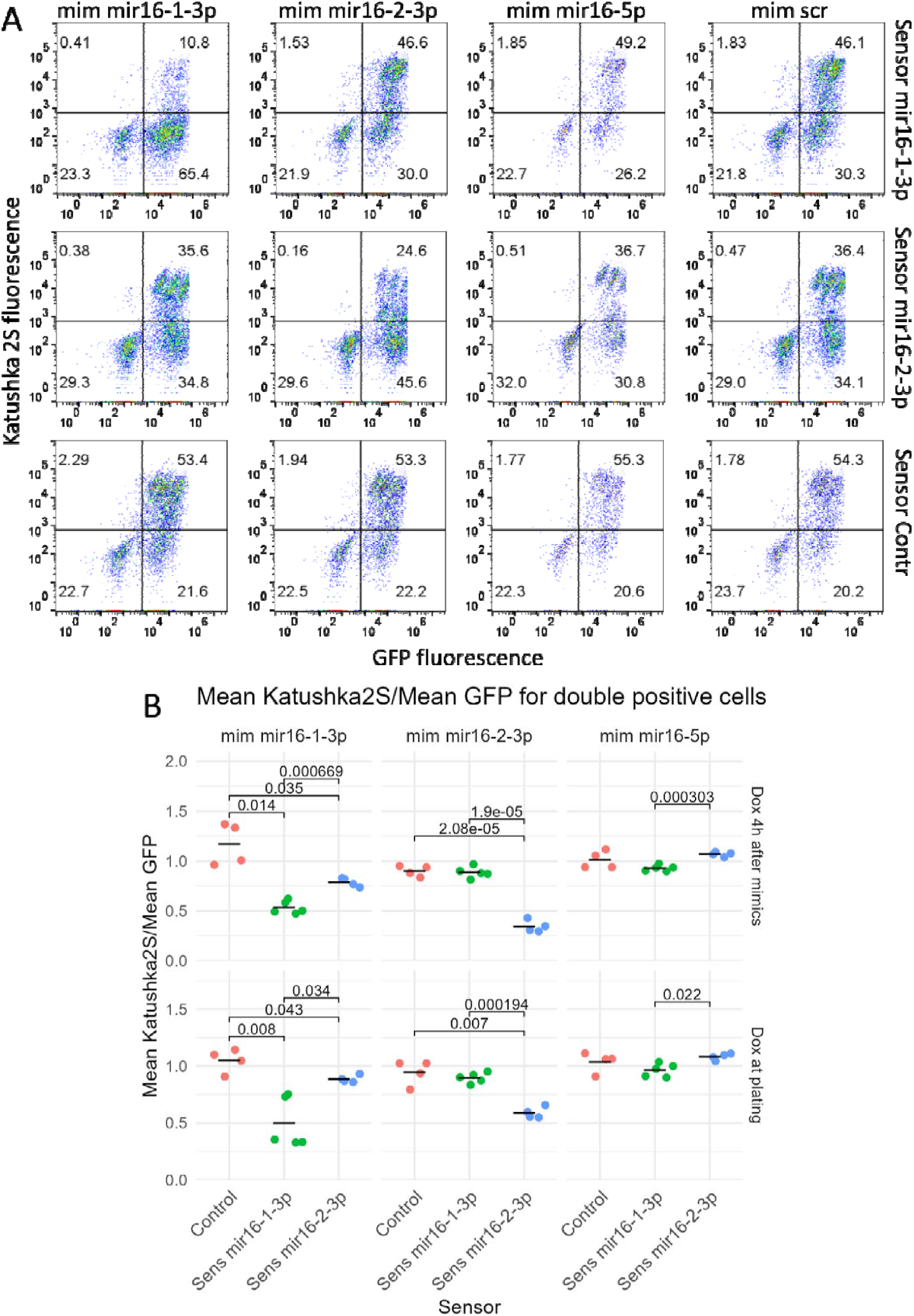
MicroRNA mimics inhibit Katushka2S fluorescence in the corresponding sensors. A. Fluorescence of individual cells after 36 hours of doxycycline induction and 40 hours post-transfection with mimics. B. Averaged normalized fluorescence of Katushka2S and GFP for cells after 36 hours of doxycycline induction and 40 hours post-transfection with mimics, as well as for cells induced by doxycycline at seeding. To make different experiments comparable, the data was normalised to the average fluorescence of the sensors after transfection with mim Scr. Significant differences are indicated using adjusted p-values.

Transfection with small RNA mimics reduced the average fluorescence of cells co-expressing Katushka and GFP while increasing the percentage of cells expressing GFP but not Katushka 2S for the respective sensors mir16-1 or mir16-2, but not for the control sensor. Additionally, the mimics decreased the fluorescence of the corresponding sensors relative to the non-specific RNA.

Changes in doxycycline concentration demonstrated that the proportion of GFP-positive cells increased from 68% to 83% as the concentration rose from 2×10^−5^ g/L to 0.005 g/L, remaining unchanged at higher, up to 0.06 g/L concentrations. Meanwhile, the proportion of cells positive only for GFP remained constant at approximately 25-30% across different experiments (Figure 3 A, B). The determination of fluorescence kinetics using live microscopy at different concentrations of doxycycline revealed that a sharp decrease in the fluorescence at concentrations below 0.0044 g/L, which corresponds to the flow cytometry data (Figure 3C).

**Figure 3.**
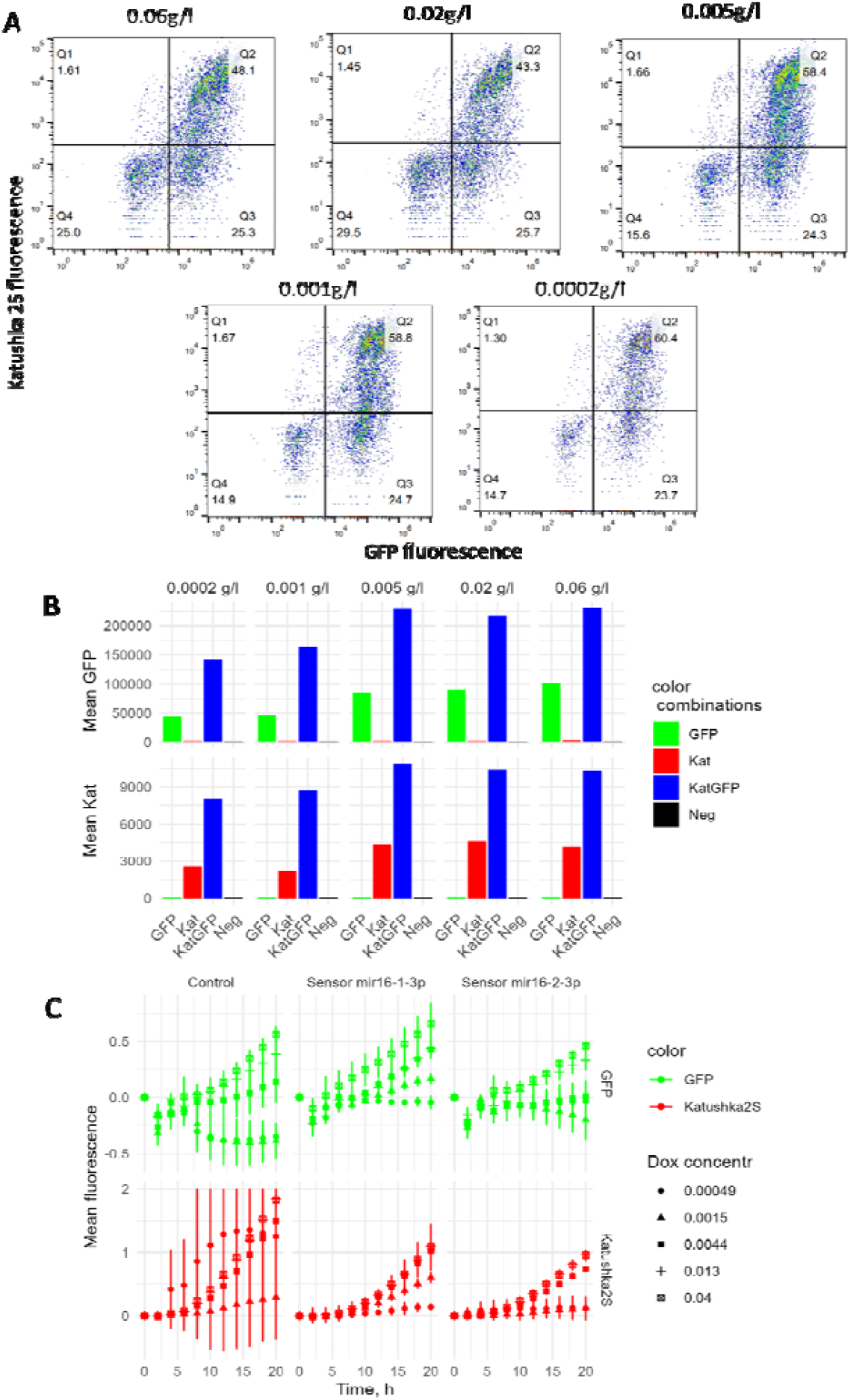
Changes of the GFP and Katushka2S fluorescence at different concentrations of doxycycline. A. THe number of GFP and Katushka positive cells increases upon the increase of doxycycline concentration. B. Averaged GFP and Katushka2S fluorescence for each quadrant, marked by the fluorofor color combinations. C. Change in the fluorescence over time for various doxycycline concentrations based on the live microscopy data.

### Kinetic measurements of the fluorescent protein synthesis facilitate determination of the mir-16-13p and mir16-2-3p miR RNA activity

Kinetic measurements of fluorescence using the JuliaStage microscope showed that cell fluorescence was detectable six hours after induction (Figure 4). In the experiments where transcription was induced at plating, we observed continued growth of fluorescence in the control samples that were not transfected with corresponding miRNA mimics (Figure 4). We observed significant differences (p.adj<0.01) in between scrambled and mir16-1-3p or mir16-2-3p mimics after only 2 or 4 hours of measurements. In contrast, in the samples that were induced by doxycycline 4h after mimic transfection, it took 8 and 10 hours for mir16-1-3p and mir16-2-3p mimics to reach a significant difference from the scrambled.

**Figure 4.**
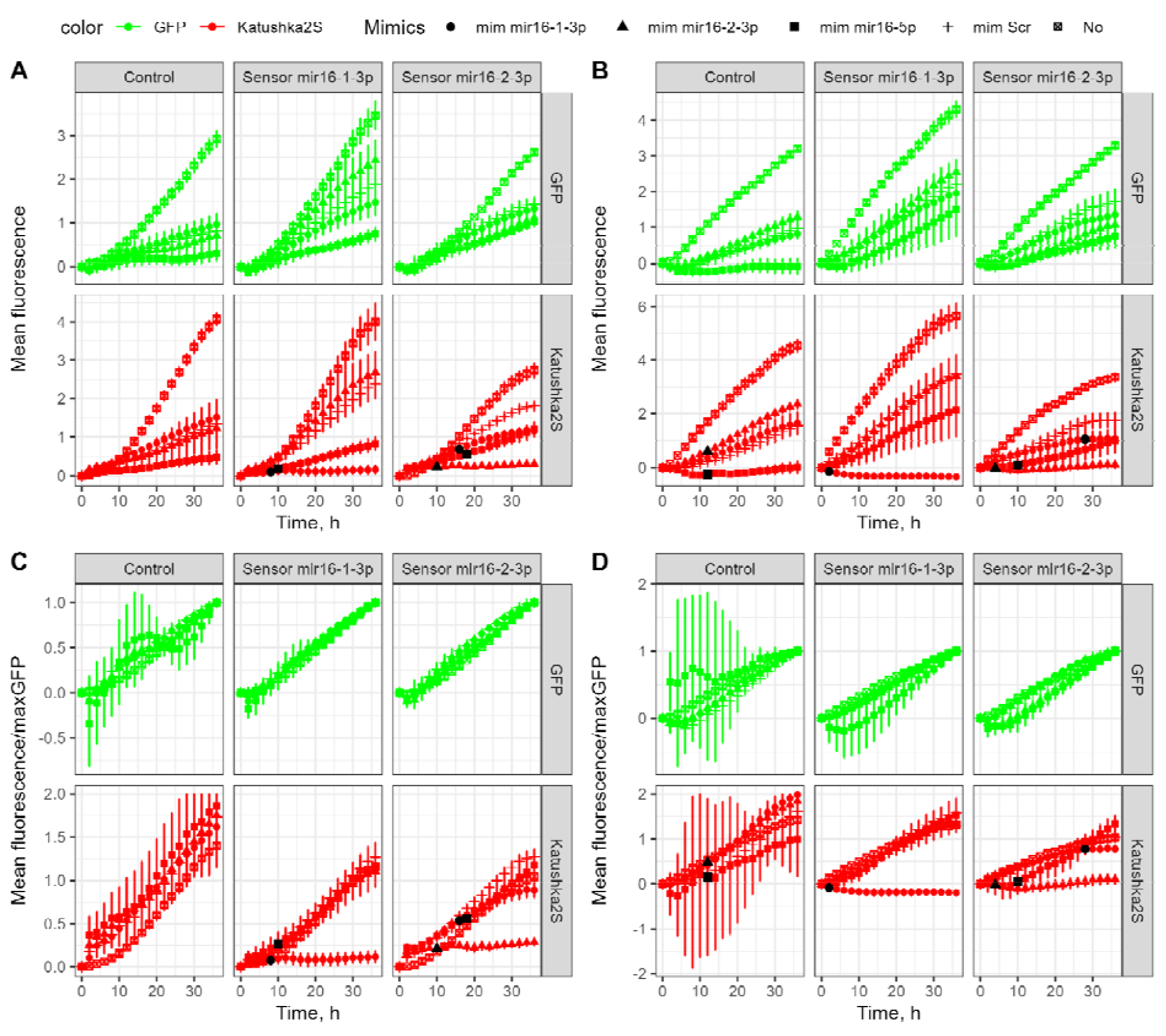
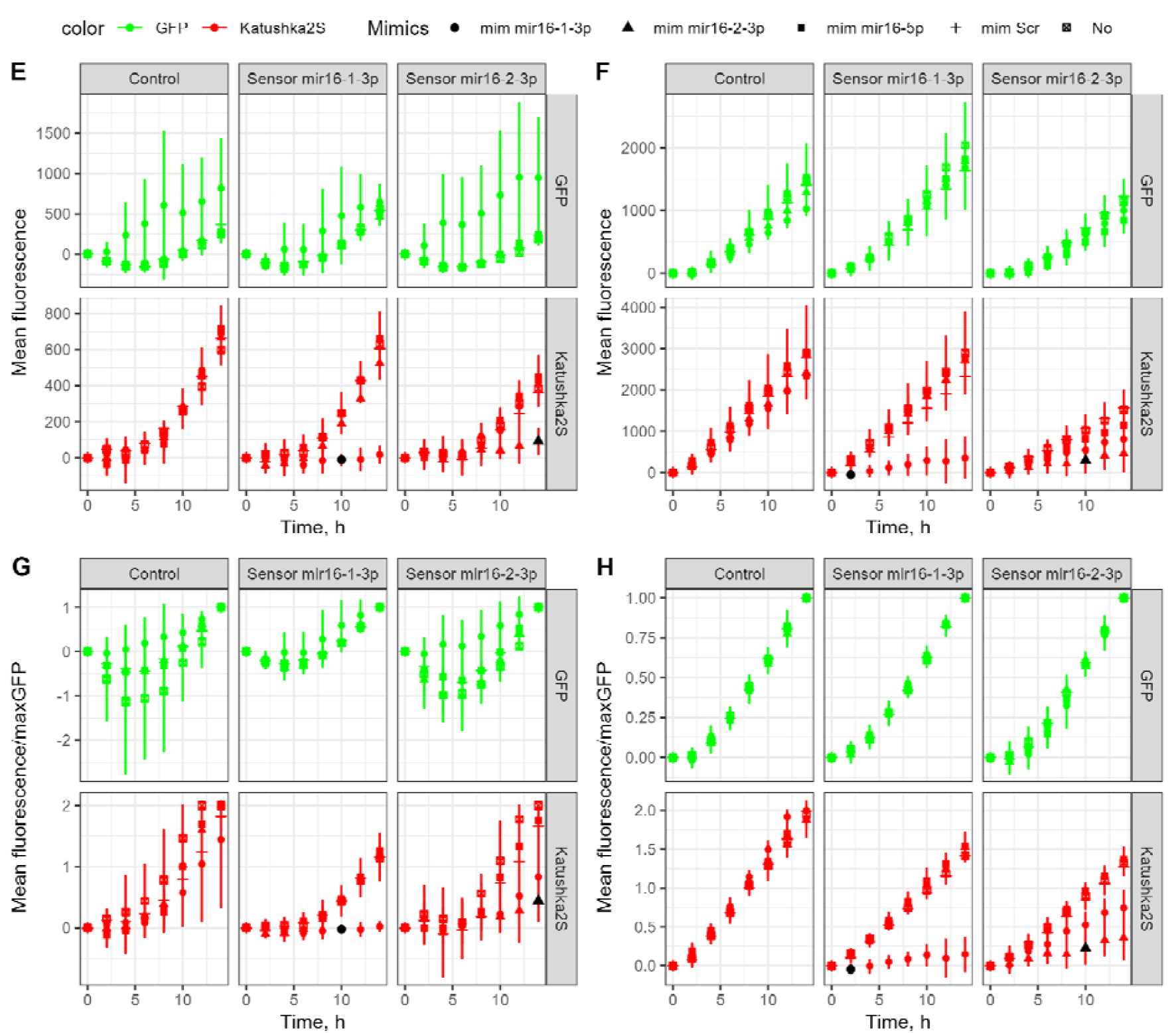
Kinetic measurements of the fluorescence and normalization to the GFP accumulated at the end of the measurements for two experiments. Black dots indicate the minimum time at which significant differences from the mim Scr (p.adj<0.01) are observed. A, B. Curves showing changes in fluorescence for doxycycline treatment started four hours after mimetic transduction (A) and at cell plating (B). C, D. The same, but the fluorescence is normalized to the amount of GFP at the end of the experiment. And for the second experiment. E, F. Curves showing changes in fluorescence for doxycycline treatment started four hours after mimetic transduction (E) and at cell plating (F). G, H. The same, but the fluorescence is normalized to the amount of GFP at the end of the experiment. Note reduction of the systematic errors related to the reduced cell viability in the first experiment.

Interestingly, in some experiments, in the Katushka2S channel values were consistent in between samples that were not transfected with corresponding miRNA mimic (Figure 4 E-F). In the first experiment, pronounced toxicity from the transfection, especially mir16-5p mimic, was observed. Consequently, for this sample, the overall fluorescence in both channels was significantly decreased (Figure 4 A-B). Normalization to the maximum value at the end of the measurements in the GFP channel showed that fluorescence in control samples and with nonspecific mimics changed proportionally, while fluorescence in the presence of mimetics complementary to the sensors decreased. Noticeably, this normalization reveled that mimic mir16-1 had some, yet not significant effect on the mir16-2-3p sensor activity (Figure 5 G-H), which was not reproduced in the initial experiment (Figure 5 A-D).

## Discussion

The final objective of these sensors is to evaluate the delivery of small RNA mimetics to tumors in vivo. Therefore, we used far-red fluorescent protein Katushka2S that was specifically designed for these purposes (34). Although, GFP has been used for fluorescence detection not only in vitro but also in vivo within tumor models (36,37), to minimize background fluorescence, particularly for in vivo applications, it is feasible to substitute the EGFP-Katushka2S pair with an alternative fluorescent protein pair, such as EYFP-Katushka2S (38,39). In addition, for in vitro applications, for simultaneous measurements using green reporters in vitro, such as green-labeled RNA mimetics, it is possible to change EGFP to EBFP-Katushka2S.

Our findings indicate that the tetracycline-induced sensor demonstrates sensitivity to exogenous mimetics mir16-1-3p and mir16-2-3p while showing insensitivity to control small RNAs. By performing a simple subtraction of the initial fluorescence in the RFP channel and normalizing it to the amount of GFP at the end of the experiment, we achieved reproducible kinetics for control samples and distinct kinetics in the presence of complementary small RNA mimetic sensors. We evaluated two methodologies for measurement. The first method entails delivering small RNA mimics followed by the induction of the reporter with doxycycline. The second method involves inducing the reporter at the time of cell seeding, with subsequent delivery of the mimics 20 hours later.

Our data suggest that both methodologies are sensitive to the delivery of the exogenous mimics with pre-induced reporter archiving differences between complementary miRNA mimics and the control mimics faster.

The vast majority of studies model siRNA action against constitutively expressed genes. In contrast, we tried to model how gene induction responds to miRNA. In the first scenario, the reporter and likely other protein response is determined by the rate of protein degradation, which is in the range of 1.5-2h for destabilised fluorescent proteins (29)(40).

The rate of the reporter response was in the range of 5-7h, which is a convolution of the miRNA synthesis and maturation and reporter protein degradation (29). In our initial experiments we failed to destabilise Katushka2S by the best modified PEST sequence (40). In contrast, protein folding is much faster, within 14 minutes for Katushka2S (34) and the decrease of protein synthesis in the similar system in response to reporter induction can be detected within 2h, similar to our results (33). In that paper, bidirectional tet-On reporter has been used to drive the expression of the renilla and firefly luciferase pair (33). Accordingly, tet-On system induction with fluorescent proteins in our paper overcomes the influence of long fluorescent protein degradation on the reporter response time. In addition, in contrast to lyciferase reporter assay, the readout can be done automatically in life cells using time-lapse microscopy at cellular resolution.

The ranges of the reporter sensitivity and specificity for the two measurement methods with pre-induced and post-induced reporter described in this study are yet to be determined.

Investigations into the reporter’s sensitivity to varying concentrations of doxycycline revealed On-Off kinetics; specifically, at doxycycline concentrations below 0.005 g/L, the reporter was either turned off or, in some instances, remained constitutively active, indicating a necessity for higher concentrations of doxycycline for reproducible measurements. However, due to potential degradation of doxycycline, we recommend establishing a concentration dependence for each antibiotic batch.

A limitation of the constructed reporter is the induction of GFP in a subset of cells, as indicated by flow cytometry analyses. One plausible explanation for this observation is that the tetracycline promoter lacks symmetry, resulting in initial GFP induction followed by Katushka2S expression.

In conclusion, we have successfully developed, for the first time, a tetracycline-induced fluorescent reporter that is sensitive to the levels of mir16-1-3p and mir16-2-3p. The modular architecture of this reporter permits further modification for the detection or simultaneous induced expression of other small RNAs or proteins. Our results demonstrate that employing kinetic measurements based on the synthesis rather than degradation kinetics of the fluorescent proteins facilitate rapid assessment of the small RNA activity.

## Acknowledgements

Pestov Nikolai Borisovich for doxycycline sample. Vadim Maximov for fruitful discussions. Victor Tatarsky for plasmid.

This work was supported by the Russian Science Foundation (project No. 23-14-00220). Bibliography

